# Model Reduction Tools For Phenomenological Modeling of Input-Controlled Biological Circuits

**DOI:** 10.1101/2020.02.15.950840

**Authors:** Ayush Pandey, Richard M. Murray

## Abstract

We present a Python-based software package to automatically obtain phenomenological models of input-controlled synthetic biological circuits from descriptive models. From the parts and mechanism description of a synthetic biological circuit, it is easy to obtain a chemical reaction model of the circuit under the assumptions of mass-action kinetics using various existing tools. However, using these models to guide design decisions during an experiment is difficult due to a large number of reaction rate parameters and species in the model. Hence, phenomenological models are often developed that describe the effective relationships among the circuit inputs, outputs, and only the key states and parameters. In this paper, we present an algorithm to obtain these phenomenological models in an automated manner using a Python package for circuits with inputs that control the desired outputs. This model reduction approach combines the common assumptions of time-scale separation, conservation laws, and species’ abundance to obtain the reduced models that can be used for design of synthetic biological circuits. We consider an example of a simple gene expression circuit and another example of a layered genetic feedback control circuit to demonstrate the use of the model reduction procedure.

## I. Introduction

Model reduction is a widely used tool in engineering design and analysis. Abstracting away the details of a system model to focus on modeling the properties of interest and its interactions is an important insight that is commonly used in control systems’ design. Reduced models are also useful to specify the desired objectives or the performance specifications of a system. To meet these objectives, the designer needs to map these reduced models to the level of system design and also mathematically characterize this mapping in order to understand and analyze the system performance. For biological systems, this is a challenge that obscures the use of mathematical models in experimental designs and analysis to some extent. One of the most common assumptions in biological model reduction is that of time-scale separation wherein processes work at different time scales. This assumption has been used to obtain reduced order models by assuming species at quasi-steady state.

A recent review paper [1] discusses basic ideas around quasi-steady state approximation (QSSA) based model reduction approaches. The paper also discusses other common approaches of model reduction such as finding slow invariant manifolds [2], computational [3] and geometric [4] singular perturbation theory in reaction kinetics models. The latter approaches are based on utilizing time-scale separation in system dynamics. However, a common downside to singular perturbation theory [5] based model reduction approaches is that finding the small parameter that transforms the system of ordinary differential equations into the required structure of singularly perturbed dynamics is non-trivial and hard to determine for general system dynamics. In [6], a method to determine the small parameters is presented and applied to the enzymatic reaction system. However, it is not yet clear how this approach would scale up to the commonly encountered models in synthetic biology. Hence, the relaxed version of singular perturbation based model reduction using QSSA [7]–[10] is more commonly used.

With standard QSSA based model reduction we do not get bounds on the error in reduction or any other form of guarantee that ensures that the reduced models are correct. The assumptions that drive the QSSA methods are usually *ad hoc* and based on understanding of the system in consideration at particular operating conditions. Due to these disadvantages, the approach cannot be systematically used in a generalized fashion or be guaranteed to work for systems under various conditions often leading to incorrect models [11, Ch. 3]. A computational approach to improve QSSA based model reduction that provides bounds on error in reduction was shown in [12]. In [13], this approach was extended to ensure that the reduced models obtained by repeated application of combinatorial QSSA leads to models that are robust with respect to parameter variations along with minimizing the errors. Our work in this paper develops further on this work.

A model reduction approach for controlled biochemical reaction systems is presented in [14]. Algebraic approaches such as state elimination using conservation laws, parameter lumping and non-dimensionalization are combined with a balanced truncation method that retains the input-output mapping for the controlled biochemical system. Balanced truncation [15]–[17] is a model reduction method that is based on proposing a transformation of system coordinates so that the unobservable and uncontrollable modes can be eliminated. These methods have been shown to be successful for linear dynamical systems [18]. However, extensions of these methods to nonlinear dynamics is still an unsolved research question. Other approaches also use transformation of coordinates to reduce a system model such as in [19] where it is shown that coordinate transformations can be proposed to bring the system into the desired singular perturbation form in order to identify the slow and the fast dynamics. Based on this approach, an automated algorithm has also been presented in [20]. However, this algorithm assumes that fast and slow species have already been recognized in the model, which is not always an easy task. To develop better and more suitable model reduction methods for synthetic biological circuits, it is clear that we would need to combine algebraic approaches that take advantage of the structure of chemical reaction level models with the general approaches that exploit time-scale separation assumptions. The results in [1] reinforce this point. Our approach in this paper is a step in this direction.

### A. Summary of results in this paper

In [13], it was shown that for an autonomous nonlinear dynamical system reduced models can be obtained in an automated fashion by choosing the set of states to collapse in the QSSA procedure by using two metrics as discussed in the later sections. We extend these results in this paper by providing results that enable us to obtain robust reduced models for general nonlinear dynamical systems that are driven by a set of inputs as well as different initial conditions. We combine this extension with the assumptions of conservation laws and species abundance to obtain phenomenological Hill function models for synthetic biological circuits. We also present a Python package that implements this algorithm. We demonstrate our method using different examples of synthetic biological circuits by obtaining commonly encountered reduced order models in an automated manner.

### B. Significance of the results

In addition to guiding the design of synthetic biological circuits, the reduced models play an important role in the analysis of the circuits using the experimentally obtained data. Since the model parameters that lead to the desired outputs are not unique for CRN level models, it can be challenging to identify all of the parameters. This issue of parameter non-identifiability has been explored in detail in the literature [21]. Using the approach in this paper, we obtain reduced order models with lower number of parameters hence helping in the analysis of these circuits. The approach can also be used to find out a mathematical description of the non-identifiable manifold as it can give expressions for the lumped parameters which correspond to this manifold.

## II. Preliminaries

We denote an eigenvalue of a matrix *P* by *λ*(*P*). The maximum eigenvalue will be denoted by *λ*_max_(*P*). For a state-dependent matrix *P* (*x*) we denote the maximum eigenvalue of *P* over all values of *x* by 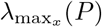. Throughout this paper, we consider the Euclidean 2-norm for vectors. For example, we use the notation *‖x ‖* for the 2-norm of *x* ∈ ℝ^*n*^ and similarly for matrices *‖ · ‖* represents the induced 2-norm. The 2-norm for a signal *x*(*t*) defined for all *t ≥* 0 is given by

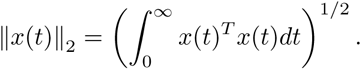

### Definition 1.

A *nonlinear dynamical system* with affine inputs is defined as

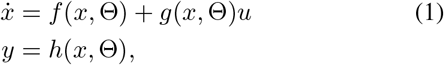

where *x* ∈ ℝ^*n*^ are the states of the system, *u* ∈ ℝ^*q*^ are the inputs and *y* ∈ ℝ^*p*^ are the outputs. The vector of model parameters are given by Θ and the respective dynamics are given by the nonlinear functions *f, g*, and *h*. We assume that the initial conditions for the system are given by *x*(0) = *x*_0_ *∈* ℝ^*n*^.

### Definition 2.

A *reduced order nonlinear dynamical system* for the system in (1) is defined as

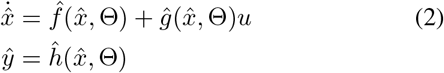

where 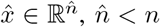 and *ŷ* ∈ ℝ^*p*^, since we assume that the number of outputs are not reduced. The initial condition for the reduced model is given by 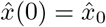.

### Definition 3.

The *sensitivity coefficient* of a state variable *x* with respect to a parameter *θ* ∈ Θ is defined as

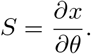

From [22], for dynamics of *x* given by the nonlinear function *f*, i.e., 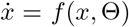 we know that the sensitivity coefficients at *x*(*t*) = *x*^*∗*^ satisfy the sensitivity equation 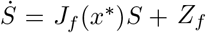, where *J*_*f*_ (*x*^*∗*^) ℝ^*n×n*^ is defined as the *Jacobian* of the nonlinear function *f* at the point *x*^*∗*^,

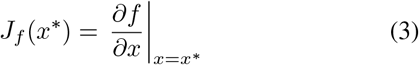

and *sensitivity to parameter* matrix for the function *f* is defined as *Z*_*f*_ *∈* ℝ^*n×*1^ and given by,

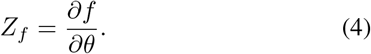

For simplicity, we often drop the subscript *f* to write *J*_*f*_ = *J* and *Z*_*f*_ = *Z*.

For simpler notations when working with the full and reduced model together, we define an augmented state vector 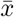 as follows

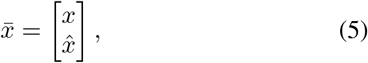

where *x* is the full state vector and 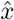 is the reduced state vector. We extend this notation to all other corresponding elements such as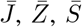, etc. as given below

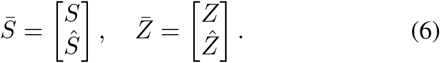

## III. Results

Before presenting the main results, it is important to note the following assumptions that we make for the rest of this paper:

1. We assume mass-action kinetics for chemical reaction network dynamics that describe the biological processes. Hence, our circuit models will be given by deterministic ordinary differential equations.
2. As discussed in the Introduction, we are only interested in the structured model reduction problem where the states of the full model retain their definitions in the reduced models. Hence, we do not consider coordinate transformations to reduce the models. As a result, the reduced state variables are related to the full state variables as follows:

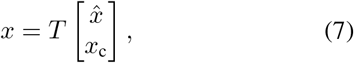

where *T* is a permutation matrix with only one nonzero element in each row or column, and *x*_c_ is the set of collapsed state variables, i.e. the variables for which dynamics are collapsed in the process of reducing the model.
3. From the previous assumption, it is also implied that the inputs and outputs will always be retained in a reduced model.
4. As given in the nonlinear dynamical system in equation (1), we assume affine input dynamics.

We are now ready to present our main results. Consider a forced nonlinear dynamical system with an input *u* given by

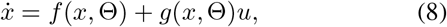

a linear relationship for the output,

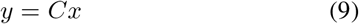

and the reduced model

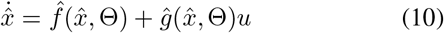

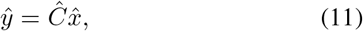

where *x* ∈ ℝ ^*n*^ and the reduced state variables 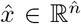 such that 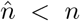. For simplicity of exposition, we assume that the input is scalar. The results that follow can be derived for systems with multiple inputs as well but with more complicated algebra. To choose the correct reduced model obtained after making the appropriate assumptions of time-scale separation, conservation laws, and species abundance assumptions, we consider the following metrics. We show how we can computationally associate these metrics with every possible reduced model to decide the best reduced model for a given system dynamics. Note that these results extend the results in [13] for non-autonomous nonlinear dynamical systems.

### A. Error metric

Similar to [13], we can derive a bound on norm of the error in the outputs *e* = ‖*y* −*ŷ* ‖_2_. Clearly, this is the most important metric of comparison between a full model and a reduced model as it compares the outputs of the two models for all time. Define an augmented system with states of the full and the reduced model stacked together as follows,

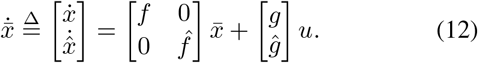

Note that we omitted the variables in the notation above for *f* (*x*, Θ), *g*(*x*, Θ), and so on for brevity. The advantage of this augmented state dynamics is that the error in the outputs *e* naturally appears as the output for this augmented system since,

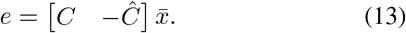

As pointed out in [12], finding a bound on the error for general nonlinear systems might not always be possible and is usually associated with finding a Lyapunov function for the nonlinear system [23]. So, in the next theorem we present a bound on the error for the linearized dynamics of the full and reduced order model and discuss the methods available for nonlinear systems next. Define,

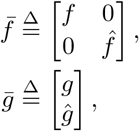

and its linearization around a point *x*^*∗*^ ∈ ℝ^*n*^, given by the Jacobian evaluated at this point,

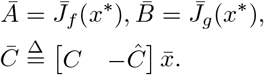

#### Theorem 1.

*If the linearized dynamics of the full and the reduced model are asymptotically stable, then the norm of the error in outputs is bounded above by*

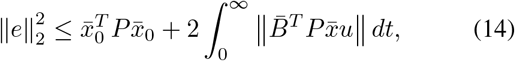

*if there exists a P* = *P*^*T*^ ≻ 0 *that satisfies* 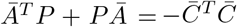.

Before proving this theorem, note that the matrix *P* can be obtained for the linearized dynamics by solving the Lyapunov equation 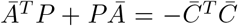 where *Ā* is the linearization of 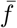 around a point in state-space. To get a bound on the error without linearizing the dynamics, we could either compute the norm of the error in a brute force way computationally or find a storage function using techniques such as SOSTOOLS [24] to find an appropriate upper bound, as shown in [12]. Also note that bound in the theorem can be further simplified as shown in [12], [13] to only depend on the reduced variable 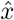 and the collapsed state variables.

*Proof*. Consider a positive definite function 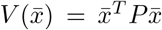 such that *P* satisfies the conditions in the theorem statement. Taking the derivative with respect to time, we get

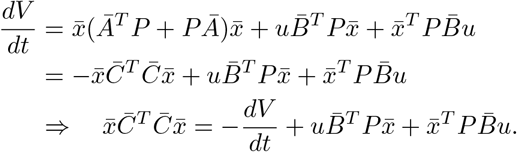

Now, for the norm of the error we can write,

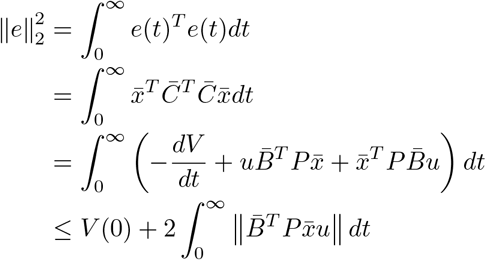

Writing 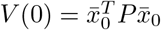 proves the theorem. □

### B. Robustness metric

In this section we discuss a metric that gives a measure of robustness of the reduced model with respect to the full model. We can use this metric to compare various possible reduced order models to find out the models for which the error in model reduction is the least sensitive to the model parameters. This metric is given by

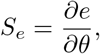

for *θ* ∈ Θ. The next result gives a bound on the robustness metric for the forced nonlinear dynamical system in equation (8).

#### Theorem 2.

*For the nonlinear controlled dynamics (1) and its reduced model (2), the norm of the sensitivity of the error in model reduction S*_*e*_ *is bounded above by*,

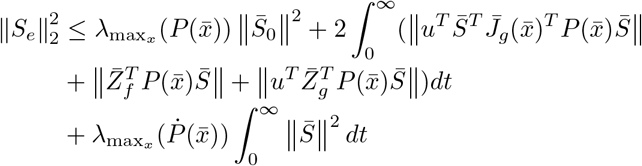

*if there exists* 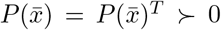 *such that* 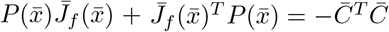 *at every point* 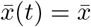 *in the system trajectory*.

Similar to the previous theorem, the condition can be simplified further as shown in [13] to only depend on the reduced variable 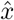 and the collapsed variable *x*_c_. This simplification follows by defining a permutation matrix *T* that separates the state variables *x* into 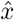 and *x*_c_ after time-scale separation assumptions have been applied. Also note that, unlike the previous theorem, it is simpler to find a matrix *P* that satisfies the given condition. This is due to the fact that the sensitivity system is a linear system where we can obtain the Jacobian matrix at each time point and use the continuous-time Lyapunov equation to solve for a *P* that satisfies the conditions required in the theorem statement.

*Proof*. The proof follows similar to the proof of the previous theorem by defining a function 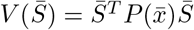 and calculating the bound for ‖*S*_*e*_ ‖. Note that 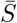 is defined according to the notation given in Section II. The additional condition on *P* helps to give the bound in the final form as given in the theorem statement. See the Appendix for full proof. □

The results above assumed that the outputs of the system are linearly related to the states *y* = *Cx*, however, we can derive similar results even without this assumption.

#### Theorem 3.

*For the system dynamics in equation (1) with output dynamics given by y* = *h*(*x*, Θ) *and the reduced model dynamics given in equation (2) with output dynamics given by* 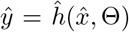, *the norm of the sensitivity of error is bounded above by*,

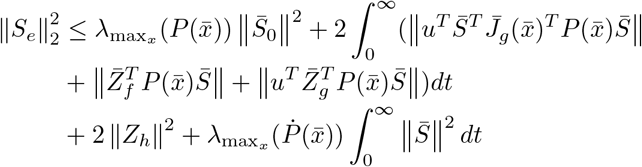

*if there exists* 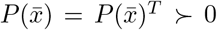 *at every point* 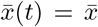 *such that* 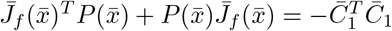, *where*

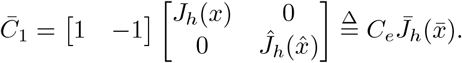

*Proof*. For *S*_*e*_, at a point 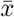, we have the following from Definition 3,

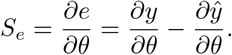

Using chain rule, we can write

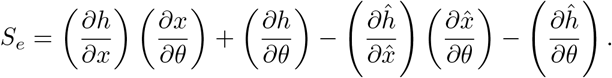

Defining the Jacobian 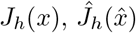 and parameter sensitivity matrices 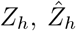 with respect to *h* and substituting back, we can write,

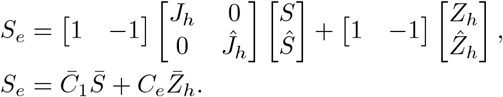

Now, consider the function 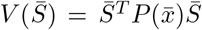 and proceed in a similar way as in the proof of previous results to write 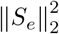,

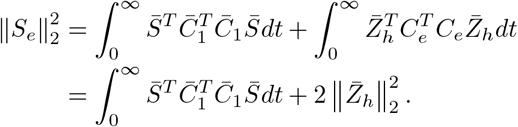

Using the result from Theorem 2 and 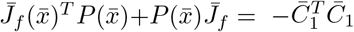,

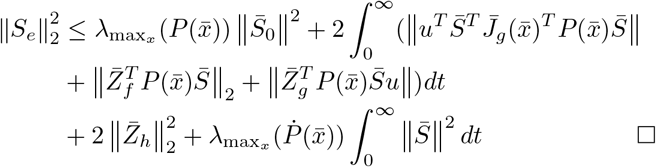

### C. Input-Output mapping

For forced nonlinear systems, in addition to the error and the robustness metric, it is also important to consider the input-output mapping so that the response of a reduced model for an input is similar to that of the full model. If the mapping from *u* ↦ *y* is linear, the computation of induced system norms is a well studied topic, see for example [25]. However, for general nonlinear systems, this is still an active research area [26] with results only available under certain structural conditions on the system dynamics. Similar to the gap-metric for linear systems [27], [28], there have been a few results on computation of the gap metric for nonlinear systems [29]. We can use similar computational results in our model reduction procedure by assigning the gap metric to each nonlinear reduced order model.

On the other hand, if we are only interested in the response to an input at steady state or at a fixed number of points in the response, then we can linearize the dynamics at these points and assess the induced system norm for each reduced model and use this as a metric while choosing a reduced order model. For the system operator *H* : *u* ↦ *y*, we define,

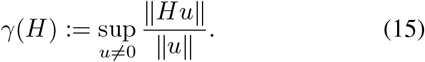

We will use this metric to demonstrate the system norm comparison between reduced models for input signals applied to the biological examples considered in the next section.

### D. Python software package

A Python based implementation of the model reduction algorithm discussed above called `AutoReduce` is available at [30]. The algorithm is similar to Algorithm 1 given in [13]. In [13], the algorithm was used to obtain a reduced order model for the circuit in [31]. We extend this algorithm using the results discussed in this paper and demonstrate its application to more general examples. The Python package works by loading an ODE model into the `System` object using Python Sympy [32]. Local sensitivity analysis using various approximate difference methods [22] and a more accurate method that solves the sensitivity system ODEs directly are also implemented in this package. With the `Reduce` class, the following important tools are available:

#### 1) Time-scale separation

To solve the time-scale separation problem for a given model, the package methods can be used to set the dynamics of the given states to collapse (*x*_c_) to zero and to automatically substitute back into the dynamics for the reduced state variables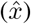. This method automates the QSSA procedure and can often be useful to compute QSSA based reduced models without specifying parameter values.

#### 2) Conservation laws

Using conservations laws, we can eliminate states that are conserved from our ODE model. We integrate this appraoch in our method similar to the method in [14]. The conservation laws can be explicitly defined and these are used in the software package accordingly to eliminate state variables and compute the reduced order models.

#### 3) Comparison metrics

The comparison metrics discussed above are implemented in this package as well. For any pair of full model and reduced model, metrics such as the norm of the error (‖*e*‖) or the robustness metric (‖*S*_*e*_‖ can be computed.

## IV. Examples

In this section, we will present biologically relevant examples to demonstrate the model reduction procedure discussed in the previous section. We start with the canonical example of the enzymatic reaction system for which we get the Michaelis-Menten kinetics as the reduced order model. We show that we can use the software package discussed above to specify the conservation laws and get the reduced order model in an automated manner.

### A. Enzymatic reaction dynamics

We create the chemical reaction network model using mass-action kinetics for the enzymatic reaction system described by the following reactions:

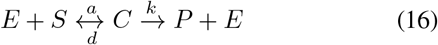

The full mass-action kinetics based model with four species is given by:

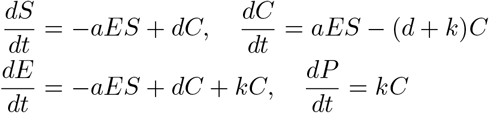

To obtain the reduced model, we use the Python package presented in this paper. After loading the `System` object in the package with the full model and the parameters given above, we specify the conservation laws for this system : *E* = *E*_*total*_ *C* and *S* + *C* + *P* = *S*_*total*_. We also specify that the output of interest for this system is *P*, the product species. The output species (P) are always retained in our model reduction procedure as discussed in the previous section. On running the software to automatically obtain the reduced order model, it first eliminates the two state-variables *E* and *S* based on the conservation laws. In the next step, it solves for time-scale separation to obtain a possible reduced order model. For this reduced model, we computed the error metric (‖*e*‖ = 9.7 ×10^−4^) as well as the robustness metric (‖*S*_*e*_ ‖) over the model initial conditions. If the metrics are lower than the desired tolerance level, then we can conclude that the final reduced model is given by:

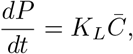

where *K*_*L*_ = *kE*_total_ is the lumped parameter and 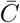 is a function of *E*_total_, *S*_total_ and model parameters given by

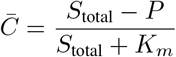

where *K*_*m*_ = (*d* + *k*)*/a*. All of the simulations and code required for computations for this example is available at [30].

### B. Gene expression : TX-TL model

Consider a model of gene expression where a gene *G* transcribes to an mRNA species *T* which translates into the protein species *X*. In this system, the concentration of *G* controls the production of the output protein *X*. Hence, the system has an input *u* = [*G*] and one output *y* = [*X*]. The model dynamics can be derived as an ODE by writing down the mass-action kinetics for these chemical reactions:

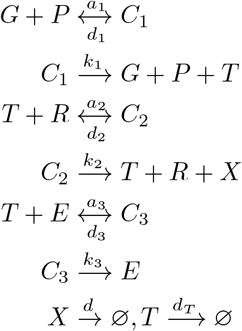

The transcription starts by the action of RNA polymerase *P* and the translation is modeled by the binding of ribosome *R* to the transcript *T*. We have also modeled the degradation of the mRNA species by the endonucleases *E*. The output species of interest in this case is the protein *X*. Since we can control the input gene concentration, we treat [*G*] as our input. The system dynamics with 8 states, *x* = [*P C*_1_ *T R C*_2_ *E C*_3_ *X*]^*T*^) and 11 parameters are given by:

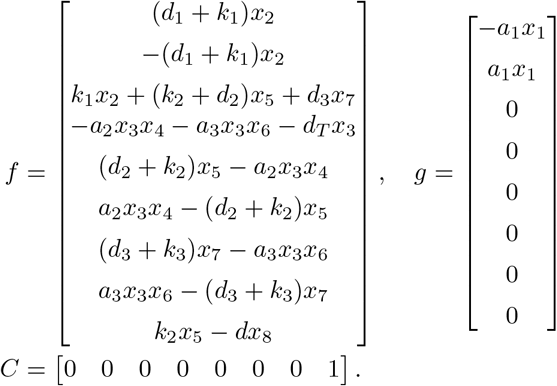

In the `AutoReduce` software package, the system model as described above can be directly loaded using symbolic variables as follows:

~~~
sys = System(x, f, g, u, C, params)
~~~

Since, we are only interested in the expression of the protein *X* driven by the input *G*, our goal is to obtain a simpler reduced order model for this system that preserves this mapping while also ensuring robust performance with respect to model parameters. We assume that the resources *P, R*, and *E* are conserved. So, we get the following conservation laws:

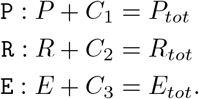

These conservation laws can be set to the `Reduce` class object created for the `System` as follows:

~~~
sys_reduce = reduce(sys)
sys_reduce.set_conservation_laws([P,R,E])
~~~

We can now obtain the various possible reduced order model for this transcription-translation model. From the dynamics of the system, we see that we would need to use the result of Theorem 2. We would also need the error metric and the metric on the input-output mapping to compare the performance of various reduced models. We can use the following utility functions available in `AutoReduce` to compute the reduced models and the related metrics:

~~~
reduced_model =
sys_reduce.solve_timescale_separation(states)
reduced_model.get_error_metric()
reduced_model.get_robustness_metric()
reduced_model.get_gamma(x_point)
~~~

In addition to these utility functions, the package also has automated functions that do all the computations required given the full model to get various possible reduced order models. For this example, we set the tolerance for maximum number of states in a reduced model to three. Among the various possible reduced models with three or fewer states, we use the computation of the metrics (as discussed above) to choose a “best” reduced order model.

#### Remark

We start by fixing the nominal parameter values and setting the initial conditions for the full CRN model. Note that the parameter values might not always be known exactly as is often the case with biological systems. However, for the numerical computations involved in the model reduction algorithm to work we need these nominal parameter values and initial conditions. The robustness metric that the algorithm computes can then be used to understand the sensitivity of each of the obtained reduced models with all the model parameters. Since we can eliminate the reduced models that have high robustness metric, the requirement of parameter values in the full model is not strict and rough guesses should suffice for the algorithm to work.

On running the algorithm, we get a set of reduced order models as shown in Figure 1. The detailed equations for some of the reduced models are given in Appendix 2. One of the reduced models that retains the states corresponding to the mRNA transcript and the output protein *X* is given below.

**Fig. 1.**
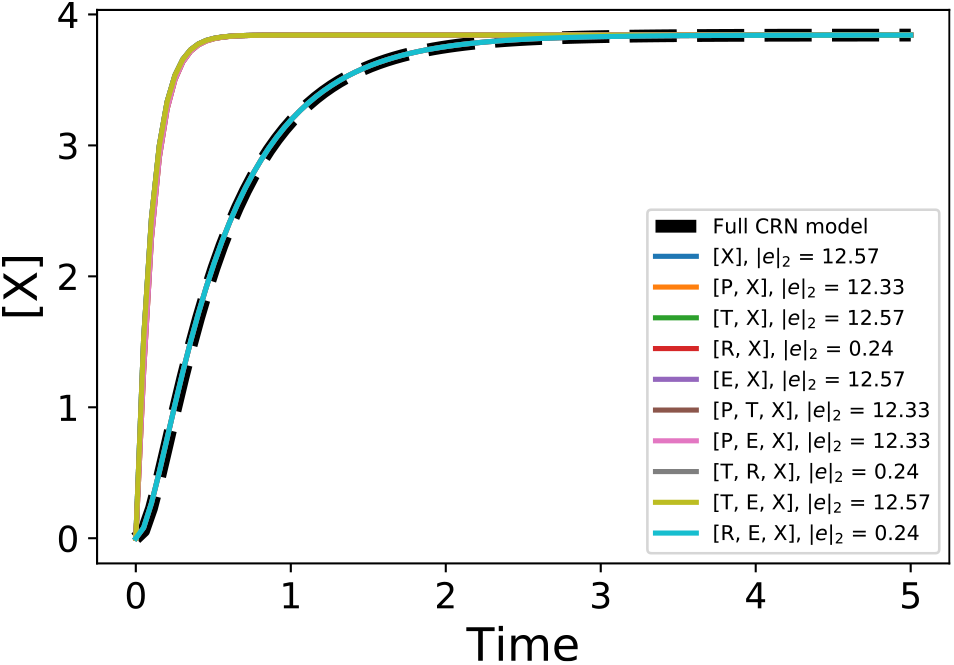
The output response of the full CRN model compared alongside the output of various reduced order models for a step input. The legend indicates each of the reduced models with the corresponding norm of the error in the output. It is clear from the figure that we cannot choose a particular reduced model based only on the error metric.

Note that this is similar to the commonly used transcription-translation models [11]. We can derive this model by lumping various parameters and making approximations based on the usual parameter values in the reduced model 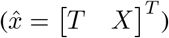 given in the Appendix 2:

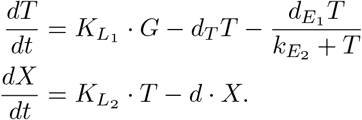

where 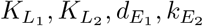 are lumped parameters. This is the best reduced model in terms of the output error metric if we wish to restrict the number of states to two. But, it is also important to check whether this reduced model preserves the input-output mapping or not and if it is robust to various model parameters. The comparison of the robustness metric for all of the reduced models is shown in the heatmap in Figure 2. Finally, in Figure 3, we show the input-output mapping for various reduced models compared to the full CRN model. As discussed in the previous section, to compute system norms it is simpler to linearize the nonlinear models. In Figure 3, the vertical lines show the points at which we linearize all of the models. Using the linearized model, we can compute any system norm of our choice. In this example, we compute the 2-norm [33, Ch.2] using the state-space matrices obtained after linearization [23] at each of the chosen point in the trajectory. Hence, based on the three metrics, we can conclude that the reduced model that retains the states *T, R*, and *X* is the best possible reduced model for this gene expression example. The model equations are given in Appendix 2. All simulations, Python package setup, and parameters for this example are available at [30].

**Fig. 2.**
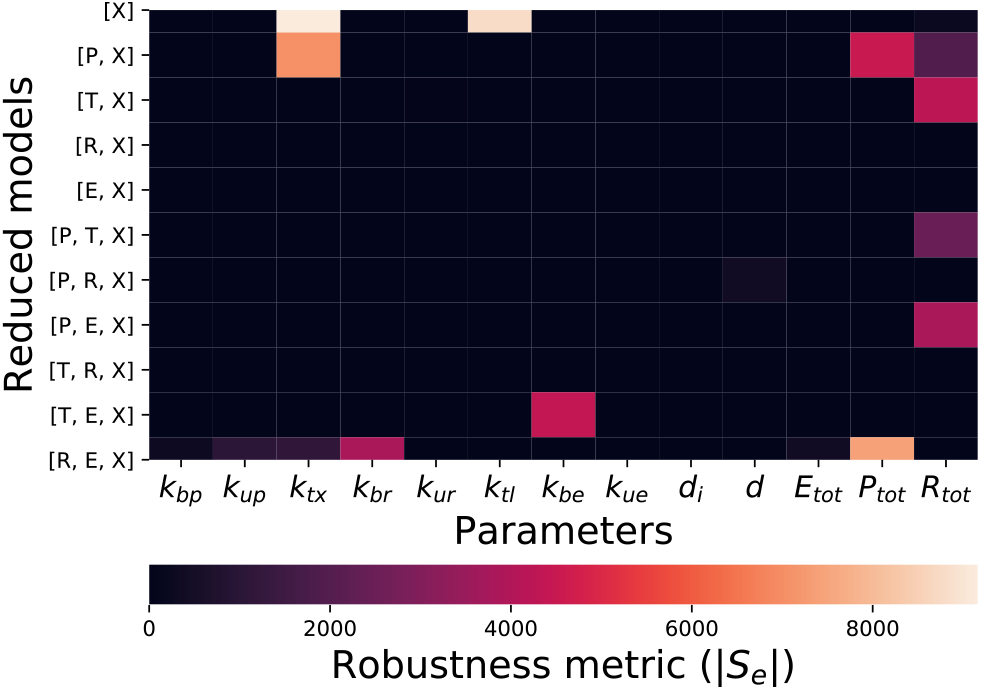
Robustness metric (‖*S*_*e*_‖) for different reduced models as computed according to the results in the previous section. From this heatmap, it is clear that to get lower sensitivity to the model parameters, we need to increase the size of 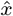 by 1 and consider models with three states retained. The reduced model with 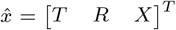 is the least sensitive to the model parameters, hence it can be a good candidate for our choice of final reduced order model.

**Fig. 3.**
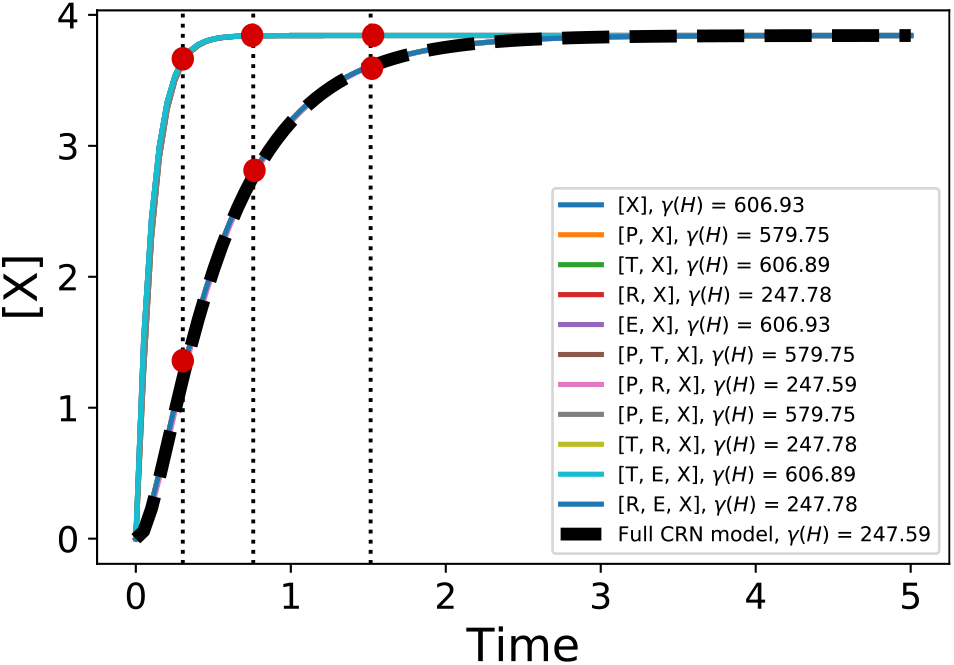
Comparison of *γ*(*H*) for each of the reduced model linearized at the points denoted. From this metric, we can conclude that the reduced model with 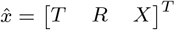 preserves the input-output mapping when compared to the full CRN model.

### C. Layered synthetic feedback controller

As the final example, we consider a genetic layered feedback controller recently shown in [34]. The circuit implements feedback control mechanisms both at the molecular and the population level to control the system performance under various disturbances. A phenomenological model was developed for this circuit and presented in [34]. This model was used for the analysis of the design and to study important properties of the system such as disturbance attenuation and stability under various perturbations. However, depending on the parts and components used to build this circuit experimentally, the detailed chemical reaction level model may or may not correspond to the phenomenological model used to make the design decisions and study performance specifications of this system. Hence, the model reduction approach presented in this paper can be used to computationally determine the phenomenological model from the chemical reaction level models in order to guide the experimental design. Since this circuit has inducer molecule concentrations that are used as inputs that control the output protein expression, the results in Theorem 1, 2, and the discussion in Section III-C are needed in order to study the reduced models.

We begin by developing a chemical reaction level model from the proposed parts and component description as shown in Figure 4:

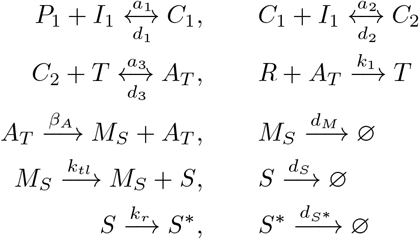

where *P*_1_ is the transcription factor Rhl-R, *I*_1_ is AHL-Rhl, *T* denotes the inactive transcript that gets activated by the dimerized complex *C*_2_ to form the active transcript *A*_*T*_. This active transcript *A*_*T*_ transcribes into the mRNA species given by *M*_*S*_. Finally, *M*_*S*_ translates into the signalling protein Cin-R denoted by *S*. A small RNA *R* acts as a regulator that represses the expression of *S* as it binds to the active transcript *A*_*T*_ to make it inactive. The protein *S* signals the production of the cis-regulatory mechanism which is governed by the following set of chemical reactions:

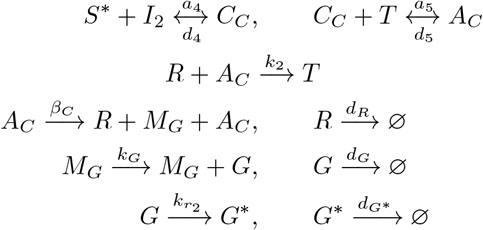

where *I*_2_ is AHL-Cin and *A*_*C*_ denotes the active transcript that leads to the downstream expression of the regulator sRNA *R. M*_*G*_ and *G* denote the mRNA and the protein species of the fluorescent protein GFP. Using mass-action kinetics, we can obtain an ODE model for the circuit. Loading this model into the software package and setting the conservation laws for this circuit, our goal is to find out reduced order models for this circuit. We use the following conservation laws:

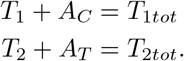

**Fig. 4.**
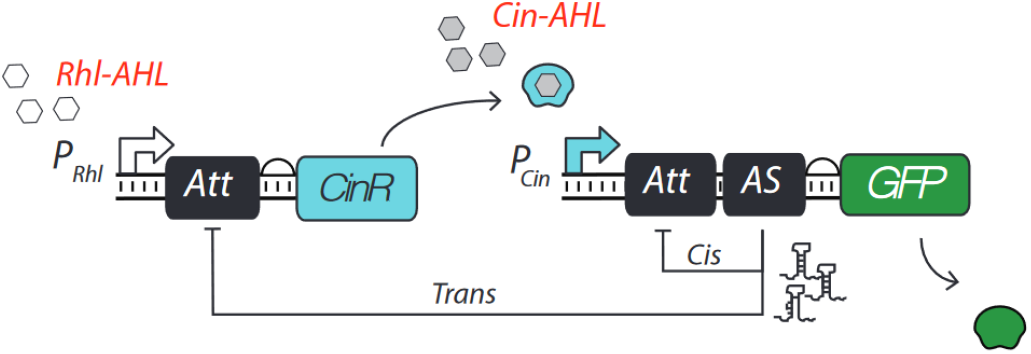
Circuit design for the layered feedback controller from [34]. In this circuit, R is an sRNA regulator that interacts with the attenuator (Att) RNA. This interaction causes transcriptional termination, and therefore down regulates the gene expression of downstream genes. The promoter P_Rhl_ is activated by the (input 1) inducer molecule (Rhl-AHL), which expresses the transcription factor CinR that activates the P_cin_ promoter in presence of the Cin-AHL inducer (input 2).

The full model after substituting the conservation laws is given by:

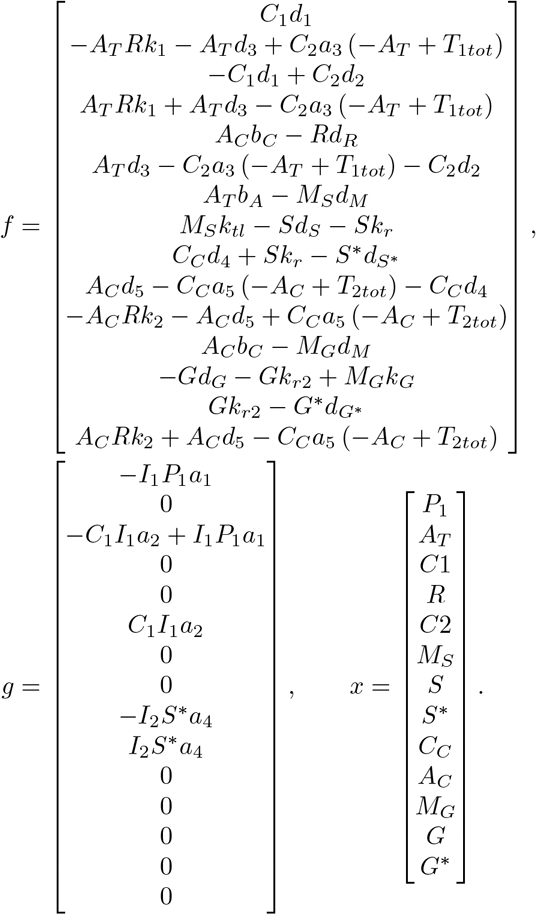

We use the `AutoReduce` package for this system model to get various reduced models. Note that we can obtain these reduced model representations without the explicit information about all the parameters. So, if we assume that the fast states are known to us *a priori*, then we can use this software package to obtain the reduced models symbolically. Using the information from [34], we can obtain a reduced model for the layered feedback control circuit. The reduced model obtained is given below. It is possible to show that this model is similar to the minimal model given in [34]. The model is obtained by retaining the states R, M_S_, A_T_, S^*∗*^, C_C_, M_G_, *G*, G^*∗*^. The reduced model is given by

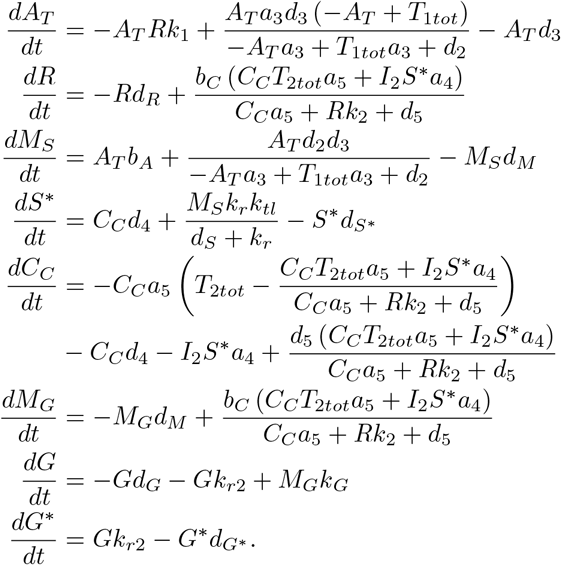

On further simplification, introducing lumped parameters, and making approximations using parameter values from [34], we can write the reduced model as follows:

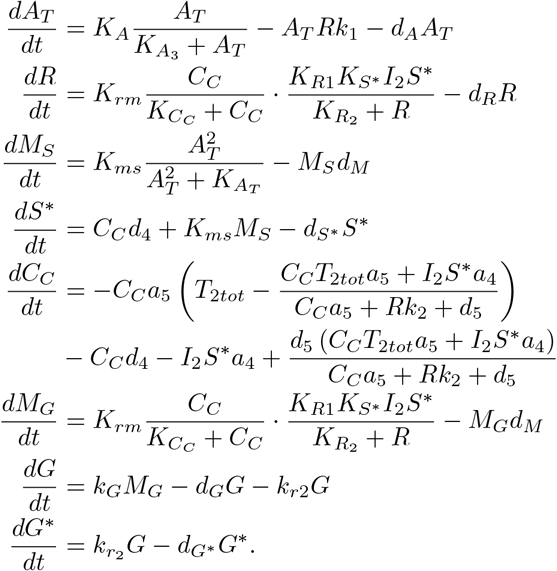

From this result, we conclude that the minimal model given in [34] is a good approximation of the full CRN level system dynamics under some assumptions. With this systematic derivation of the reduced model, we can analyse the parameter regimes in which the minimal model is close to the full model.

## V. Conclusion

Our main result in this paper gives a structured model reduction approach for forced nonlinear dynamical systems that model the chemical reactions occurring in a biological process using mass-action kinetics. We extended the results in [13] to give model reduction tools for controlled synthetic biological circuits (i.e. for systems with inputs). We derived three different metrics to compare various possible reduced order models viz. the normed error in outputs to compare the output responses between the full model and the reduced models, the sensitivity of error with respect to model parameters to compare the robustness of reduced models, and an induced norm metric to compare the input-output mapping for systems driven by inputs. Using these results, we presented an automated computational pipeline to derive phenomenological models of biological circuits directly from chemical reaction level descriptions. We implemented this automated pipeline as a Python package [30] and demonstrated its utility using examples of model reduction of a simple gene expression transcription-translation circuit model and a layered genetic feedback controller.

An important direction of future work could be to mathematically characterize the mapping from the chemical reaction level models to the phenomenological models describing the system behavior. The computational pipeline developed in this paper might be helpful towards that end.

## Acknowledgment

We would like to thank Chelsea Hu for her help with the modeling of the layered feedback controller example. This research is sponsored in part by the National Science Foundation under grant number: CBET-1903477 and the Defense Advanced Research Projects Agency (Agreement HR0011-17-2-0008). The content of the information does not necessarily reflect the position or the policy of the Government, and no official endorsement should be inferred.

## Appendix 1

*Proof*. For the nonlinear system in (8), we can write the sensitivity coefficients at any point *x*(*t*) = *x*^*∗*^ as

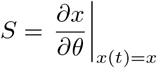

where *θ* ∈ Θ. Using chain rule, we can derive the sensitivity system equation given by

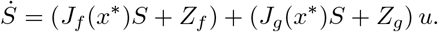

for a scalar *u*. Similarly, we can write a relation for the reduced model and hence for the augmented system given in equation (12). For the augmented system, the sensitivity coefficients are denoted by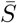. For this system, at point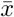, consider a function 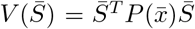 for 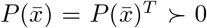 that satisfies the conditions given in the theorem statement. Taking the derivative of *V* with respect to time, we can write,

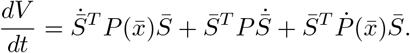

Substituting for 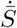 using the sensitivity system equation derived above as 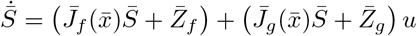 we get

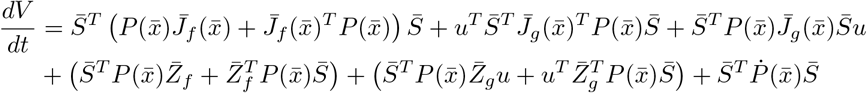

Now if there exists a 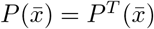 such that

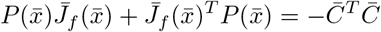

then we get the following bound by manipulating the 2-norm of *S*_*e*_,

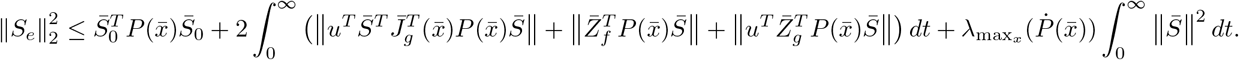

Finally, applying a bound on the initial conditions for the sensitivity coefficients, we get

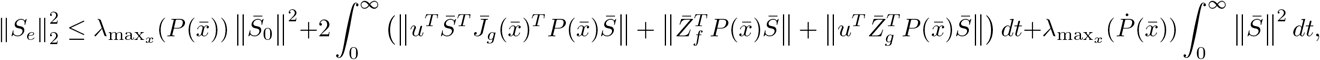

which gives us the desired result. This can be further simplified as shown in [13] to express this bound only in terms of 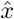, the reduced state variables and *x*_c_, the collapsed state variables. □

## Appendix 2

### Reduced models for gene expression example

The model equations for various reduced order models discussed in the Results section are given below. For 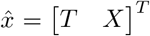we have

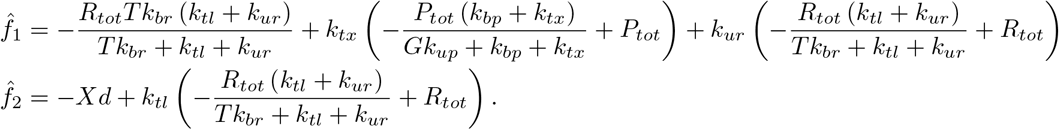

For 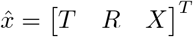,we have

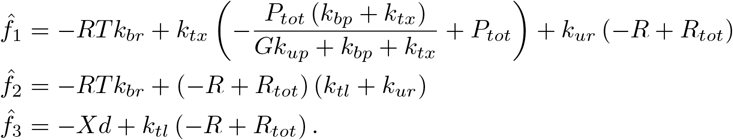

For 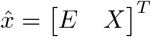we have

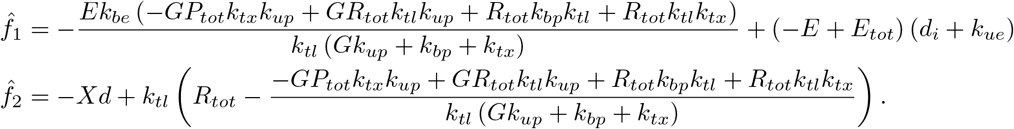

For 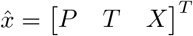we have

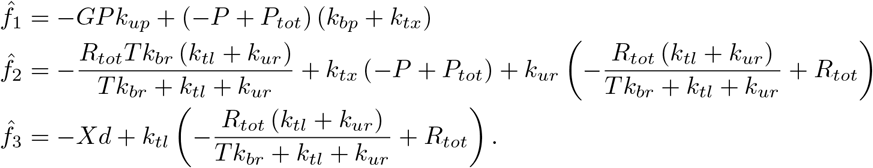

All other reduced model equations, simulations, and computations are available in [30]. For a detailed study of gene expression models, the reader is referred to our latest work in [35].

